## Supplementary material for "Variability in radiocesium activity concentration in growing hardwood shoots in Fukushima, Japan": S1 File

/*
 * Modeling of shoot 137Cs with variation in sd
 * random effects on mean but not sd
 *
 * Thanks to Mr. Matsuura (HOXO-M Inc.) for helping to make this model
 */

functions {
  real f(array[] real Y, array[] real theta, real x) {
    return normal_lpdf(x | 0, theta[2])
           + normal_lpdf(Y | theta[1] + x, theta[3]);
  }

  real log_lik_Simpson(array[] real Y, array[] real theta,
                       real x_lower, real x_upper, int M) {
    vector[M + 1] lp;
    real h;
    h = (x_upper - x_lower) / M;
    lp[1] = f(Y, theta, x_lower);
    for (m in 1:(M %/% 2))
      lp[2 * m] = log(4) + f(Y, theta, x_lower + h * (2 * m - 1));
    for (m in 1:(M %/% 2 - 1))
      lp[2 * m + 1] = log(2) + f(Y, theta, x_lower + h * 2 * m);
    lp[M + 1] = f(Y, theta, x_upper);
    return log(h / 3) + log_sum_exp(lp);
  }
}

data {
  int<lower=0> N_obs;                        // Number of observations
  int<lower=0> N_sites;                      // Number of sites
  vector<lower=0>[N_obs] Y;                  // Measured observations
  array[N_obs] int<lower=1, upper=N_sites> Site; // Site index
}

transformed data {
  vector[N_obs] logY = log(Y);
  array[N_sites, N_obs] real siteY;
  array[N_sites] int pos = rep_array(1, N_sites);
  int M = 1000;
  
  for (n in 1:N_obs) {
    siteY[Site[n], pos[Site[n]]] = logY[n];
    pos[Site[n]] += 1;
  }
}

parameters {
  real mu_bar;             // Overall mean
  vector[N_sites] eps_mu;  // Random effect on mean by site
  real<lower=0> sigma;     // Standard deviation
  real<lower=0> s1;        // Standard deviation of site ran.eff.
}

transformed parameters {
  vector[N_sites] mu = mu_bar + eps_mu;
}

model {
  for (n in 1:N_obs)
    logY[n] ~ normal(mu[Site[n]], sigma);
  eps_mu ~ normal(0, s1);

  mu_bar ~ normal(0, 10);
  sigma ~ normal(0, 5);
  s1 ~ normal(0, 5);
}

generated quantities {
  vector[N_obs] yrep;    // replicated data for PPC (log-scale)
  array[N_sites] real log_lik;

  // replicated data for posterior predictive check
  for (n in 1:N_obs)
    yrep[n] = normal_rng(mu[Site[n]], sigma);

  // log liklihood for calculation of WAIC
  {
    array[3] real theta = {mu_bar, s1, sigma};
    for (s in 1:N_sites)
      log_lik[s] = log_lik_Simpson(siteY[s, 1:(pos[s] - 1)], theta,
                                   -5 * s1, 5 * s1, M);
  }
}
