## Supplementary material for "Variability in radiocesium activity concentration in growing hardwood shoots in Fukushima, Japan": S2 File

/*
 * Generate the log likelihood to calculate WAIC
 * with a random effect on sd in additon to mean
 *
 * Thanks to Mr. Matsuura (HOXO-M Inc.) for helping to make this model
 */

functions {
  real f(array[] real z, array[] real theta, real x, real y) {
    return normal_lpdf(x | 0, theta[2])
           + normal_lpdf(y | 0, theta[4])
           + normal_lpdf(z | theta[1] + x, theta[3] * exp(y));
  }

  real log_lik_Simpson_2d(array[] real z, array[] real theta,
                          real x_lower, real x_upper, real y_lower, real y_upper, 
                          matrix W, int M) {
    matrix[M+1, M+1] lp;
    real h_x = (x_upper - x_lower) / M;
    real h_y = (y_upper - y_lower) / M;

    for (i in 1:(M + 1))
      for (j in 1:(M + 1))
        lp[i, j] = log(W[i, j])
                   + f(z, theta, x_lower + h_x * (i - 1), y_lower + h_y * (j - 1));
    return log(h_x / 3) + log(h_y / 3) + log_sum_exp(lp);
  }
}

data {
  int<lower=0> N_obs;                        // Number of observations
  int<lower=0> N_sites;                      // Number of sites
  vector<lower=0>[N_obs] Y;                  // Measured observations
  array[N_obs] int<lower=1, upper=N_sites> Site; // Site index
}

transformed data {
  vector[N_obs] logY = log(Y);
  array[N_sites, N_obs] real siteY;
  array[N_sites] int pos = rep_array(1, N_sites);
  int M = 500;
  vector[M + 1] V_Simpson;
  matrix[M + 1, M + 1] W_Simpson;
  
  for (n in 1:N_obs) {
    siteY[Site[n], pos[Site[n]]] = logY[n];
    pos[Site[n]] += 1;
  }

  V_Simpson[1] = 1;
  for (m in 1:(M %/% 2))
    V_Simpson[2 * m] = 4;
  for (m in 1:(M %/% 2 - 1))
    V_Simpson[2 * m + 1] = 2;
  V_Simpson[M + 1] = 1;
  W_Simpson = V_Simpson * V_Simpson';
}

parameters {
  real mu_bar;                  // Overall mean
  vector[N_sites] eps_mu;       // Random effect on mean by site
  real<lower=0> sigma_bar;    // Standard deviation
  vector[N_sites] eps_sigma;    // Random effect on sd by site
  real<lower=0> s1;           // Standard deviation of site ran.eff. of mean
  real<lower=0> s2;           // Standard deviation of site ran.eff. of sd
}

transformed parameters {
  vector[N_sites] mu = mu_bar + eps_mu;
  vector[N_sites] sigma = sigma_bar * exp(eps_sigma);
}

model {
  for (n in 1:N_obs)
    logY[n] ~ normal(mu[Site[n]], sigma[Site[n]]);
  eps_mu ~ normal(0, s1);
  eps_sigma ~ normal(0, s2);

  mu_bar ~ normal(0, 10);
  s1 ~ normal(0, 5);
  sigma ~ normal(0, 5);
  s2 ~ normal(0, 5);
}

generated quantities {
  vector[N_obs] yrep;    // replicated data for PPC (log-scale)
  array[N_sites] real log_lik;

  // replicated data for posterior predictive check
  for (n in 1:N_obs)
    yrep[n] = normal_rng(mu[Site[n]], sigma[Site[n]]);

  // log liklihood for calculation of WAIC
  {
    array[4] real theta = {mu_bar, s1, sigma_bar, s2};
    for (s in 1:N_sites)
      log_lik[s] = log_lik_Simpson_2d(siteY[s, 1:(pos[s] - 1)], theta,
                                      -5 * s1, 5 * s1, -5 * s2, 5 * s2,
                                      W_Simpson, M);
  }
}
