## Supplementary material for "Variability in radiocesium activity concentration in growing hardwood shoots in Fukushima, Japan": S3 File

/*
 * Modeling of shoot 137Cs with variation in sd
 */

data {
  int<lower=0> N_obs;                            // Number of observations
  int<lower=0> N_sites;                          // Number of sites
  int<lower=0> N_spc;                            // Number of species
  vector<lower=0>[N_obs] Y;                      // Measured observations
  array[N_obs] int<lower=1, upper=N_sites> Site; // Site index
  array[N_obs] int<lower=1, upper=N_spc> Spc;    // Species index
  array[N_obs] int<lower=0, upper=1> Org;        // Individual origin
                                                 // 0: sprout, 1: plantation
}

transformed data {
  vector[N_obs] logY = log(Y);
}

parameters {
  real mu_bar;                           // Overall mean
  vector[N_sites] eps_site;              // Random effect on mean by sites
  vector[N_spc] eps_spc;                 // Random effect on mean by species
  real beta;                             // Coefficient
  array[3] real<lower = 0> sigma;        // Standard deviations
}

transformed parameters {
  vector[N_obs] mu;

  for (n in 1:N_obs)
    mu[n] = mu_bar + beta * Org[n] + eps_site[Site[n]] + eps_spc[Spc[n]];
}

model {
  logY ~ normal(mu, sigma[1]);
  eps_site ~ normal(0, sigma[2]);
  eps_spc ~ normal(0, sigma[3]);

  mu_bar ~ normal(0, 10);
  beta ~ normal(0, 10);
  sigma ~ normal(0, 5);
}

generated quantities {
  vector[N_obs] yrep;    // replicated data for PPC (log-scale)
  vector[N_sites] ypred; // predicition for each site
  real log_ypred_site;   // log predicition for unobserved site
  real log_ypred_ind;    // log predicition for individual of unobserved site
  real ypred_site;       // predicition for unobserved site
  real ypred_ind;        // predicition for individual of unobserved site

  // replicated data for posterior predictive check
  for (n in 1:N_obs) {
    yrep[n] = normal_rng(mu[n], sigma[1]);
  }

  // prediction of individual 137Cs concentration for each observed site
  // without species random effect
  for (m in 1:N_sites) {
    ypred[m] = normal_rng(mu_bar + eps_site[m], sigma[1]);
    ypred[m] = exp(ypred[m]);
  }

  // prediction of site and individual 137Cs concentration
  // for spout stumps (Org = 0) in unobserved site
  // without species random effect
  {
    real new_eps_site = normal_rng(0, sigma[2]);
    log_ypred_site = mu_bar + new_eps_site;
    log_ypred_ind = normal_rng(log_ypred_site, sigma[1]);
    ypred_site = exp(log_ypred_site);
    ypred_ind = exp(log_ypred_ind);
  }
}
